# Successful regeneration of the adult zebrafish retina is dependent on inflammatory signaling

**DOI:** 10.1101/2023.08.11.552956

**Authors:** Oliver Bludau, Anke Weber, Viktoria Bosak, Veronika Kuscha, Kristin Dietrich, Stefan Hans, Michael Brand

## Abstract

Inflammation can lead to persistent and irreversible loss of retinal neurons and photoreceptors in mammalian vertebrates. In contrast, in the adult zebrafish brain, acute neural inflammation is both necessary and sufficient to stimulate regeneration of neurons. Here, we report on the critical, positive role of the immune system to support retina regeneration in adult zebrafish. After sterile, ablation of photoreceptors by phototoxicity, we find rapid response of tissue-resident microglia and neutrophils, which returns to homeostatic levels within 14 days post lesion. Pharmacological or genetic impairment of immune cell reactivity results in a reduced Müller glia stem cell response, seen as decreased reactive proliferation, and a strikingly reduced number of regenerated cells from them, including photoreceptors. Conversely, injection of the immune stimulators flagellin, zymosan, or M-CSF into the vitreous of the eye, in spite of the absence of a retinal lesion, leads to a robust proliferation response and the up-regulation of regeneration-associated marker genes in Müller glia. Our results suggest that neuroinflammation is a necessary and sufficient driver for retinal regeneration in the adult zebrafish retina.

## Introduction

Immune system activation is one of the first responses to tissue damage, e.g. by infection, disease or injury. Cells of the immune system (leukocytes) can recognize invading pathogens or factors that are secreted by damaged or dying cells (Ferrero-Miliani et al., 2007). Subsequently, leukocytes accumulate at the affected area, removing pathogens and clearing cellular debris, thus supporting the reestablishment of a physiological balance (Nathan & Ding, 2010). Conversely, if inflammation cannot be resolved, a detrimental chronic inflammation can occur, causing progressive tissue damage and pathology (Zhou et al., 2016).

In the lesioned mammalian central nervous system (CNS), accumulation of reactive astrocytes often results in a glial scar that acts as a barrier for successful regeneration (Buffo et al., 2008; Fitch & Silver, 2008; Sofroniew, 2009). Similarly, neurodegenerative diseases like Parkinson’s or Alzheimer’s show characteristics of chronic inflammation, causing subsequent neuronal death (Amor et al., 2010; Herrero et al., 2015; Kinney et al., 2018).

In contrast to mammals, lesioning of the zebrafish CNS results in a strong regenerative response despite the initial manifestation of inflammation (Kroehne et al., 2011; Kyritsis et al., 2012; Kizil et al., 2015; Bosak et al., 2018; Mitchell et al., 2018; Tsarouchas et al., 2018; White et al., 2017; Silva et al., 2020; Zhang et al., 2020). Whereas immune system activation in the mammalian CNS is typically detrimental for regeneration, studies in zebrafish demonstrated a strong beneficial link between an immune response and neural stem cell reactivity (Kyritsis et al., 2012; Aurora & Olson, 2014; Bosak et al., 2018; Tsarouchas et al., 2018). Remarkably, in the zebrafish adult telencephalon, inflammation is required to initiate a successful regenerative response, and a lipid inflammatory cue, leukotriene-C4, is sufficient to stimulate proliferation of radial glia-type stem cells (Kyritsis et al., 2012; Kizil et al., 2015).

In the retina, zebrafish Müller glial cells similarly act as stem cells, and generate neuronal precursor cells (NPCs) in response to retinal lesion (Lenkowski & Raymond, 2014; Goldman, 2014; Gorsuch & Hyde, 2014). These cells then amplify, migrate to the lesion site, and differentiate into the lost neuronal subtypes and thus gradually restore vision (Raymond et al., 2006; Hammer et al., 2022). Dying neurons release the proinflammatory cytokine TNF-α, triggering the regenerative response of Müller glia (Conner et al., 2014; Nelson et al., 2013). Likewise, the inflammation-associated factors Interleukin-11 and TGF-ß stimulate Müller glia cell cycle re-entry and NPC generation (Lenkowski et al., 2013; Zhao et al., 2014). Furthermore, microglia – the CNS tissue resident macrophages – support this initial regenerative response by secreting proinflammatory factors (Kizil et al., 2015; Conedera et al., 2019; Iribarne & Hyde, 2022; Zhang et al., 2020). However, to date the molecular pathways involved in damage recognition, stem cell proliferation, neuronal precursor cell amplification and differentiation during retinal regeneration are poorly understood; in particular, the role of inflammation is unclear (Lahne et al., 2020; Lenkowski & Raymond, 2014; Mitchell et al., 2018, 2019; Iribarne & Hyde, 2022).

Here, we analyze the contribution of inflammation to regeneration using a non-invasive, sterile phototoxic ablation model of photoreceptor cells, as the key cell type affected by retinal disease, in the adult zebrafish retina (Mitchell et al., 2018; Silva et al., 2020; Weber et al., 2013; Zhang et al., 2020). Following light lesion, we observe strong convergence of tissue resident microglia to the lesion site. Blocking inflammation pharmacologically caused reduced reactive proliferation of Müller glia stem cells and impaired photoreceptor regeneration. Similarly, in a genetic model of microglia deficiency, we find that microglia are required to support Müller cell proliferation. Conversely, when the immune stimulators flagellin, zymosan, or M-CSF are injected into the vitreous of the eye, notably in the absence of retinal lesion, Müller glial cells are triggered to undergo reactive proliferation and regeneration-associated marker gene expression. Taken together, our results show that in the regeneration-competent adult zebrafish retina, acute inflammation is an important positive regulator of retina regeneration.

## Results

### Leukocytes react to sterile ablation of photoreceptor cells

The immune system of vertebrates rapidly responds to retinal damage. In larval zebrafish, *mpeg1*:mCherry positive monocytes react to photoreceptor ablation by rapid migration towards the site of lesion (White et al., 2017). Similarly, microglia respond by accumulation and phagocytosis of debris in response to neurotoxic ablation of inner retinal cells in adult zebrafish (Mitchell et al., 2018). To investigate if leukocytes are recruited in an injury model at adult stages, we used intense diffuse light that causes sterile ablation of all photoreceptor subtypes by phototoxicity (Weber et al., 2013). In this model, due to the refractive properties of the adult zebrafish visual system, the photoreceptors in a central stripe of the retina are ablated, whereas ventral and dorsal retina is much less affected and conveniently serves as an internal control area (Weber et al., 2013; Fig. 1 D). We analyzed the accumulation and appearance of leukocytes in the central retina on sections by immunohistochemistry for L-Plastin, a pan-leukocyte marker (Redd et al., 2006; Kroehne et al. 2011) in *Tg(mpeg1:mCherry)* reporter animals, labelling the monocyte lineage, including macrophages and microglia (Figure 1A). In comparison to unlesioned (sham) controls, L-Plastin positive cells accumulated in the central lesion zone already at 2 day post lesion (dpl; Figure 1A). Whereas leukocytes showed a ramified morphology in sham control retinae, their appearance changed to an amoeboid and swollen shape at 2 dpl (insets in Figure 1A, D), as described by Mitchell et al., 2018 upon neurotoxic lesion. At 2 dpl an increased number of L-Plastin positive leukocytes at the outer nuclear layer (ONL) in comparison to sham controls is revealed (Figure 1A, B). In flatmounts of *Tg(mpeg1:mCherry) x Tg(opn1sw1:GFP)* double transgenic zebrafish that express GFP as a marker in the entire UV cone population, displaying the central lesion by the absence of GFP, the accumulation of monocytes can nicely be observed (Figure 1D). While in sham, microglia display an equal distribution and a ramified structure they accumulate in the central lesion area. Interestingly they disappear from the unlesioned peripheral regions. In a time course, the number of L-Plastin positive cells peaked at 2 dpl and subsequently declined to sham levels at 14 dpl (Figure 1B). To further determine the identity of retinal leukocytes, we analyzed immunoreactivity of L-Plastin in transgenic *Tg(mpeg1:mCherry)* animals.In sham control retinae, *mpeg1*:mCherry positive cells always co-expressed L-Plastin, indicating that all tissue resident homeostatic leukocytes were of the monocyte lineage, namely microglia (Figure 1A, S1A). In contrast, additional L-Plastin positive, but *mpeg1*:mCherry negative cells could be detected upon injury at 2 dpl (insets in Figure 1A). Subsequent quantification of the number of L-Plastin and *mpeg1*:mCherry double positive cells revealed a decrease from 99% to approximately 83%, indicating the presence of another cell type of the leukocyte lineage at the lesion site (Figure S1A). Neutrophils are known to rapidly respond to tissue damage and are labeled by transgenic *Tg(mpo:GFP)* (Renshaw et al., 2016; Kurimoto et al., 2013; Figure 1D, S1B). In contrast to sham controls, which never showed any *mpo*:GFP positive cells in the homeostatic retina, we observed rapid accumulation of neutrophils after light lesion. *mpo*:GFP positive cells first appeared in the retina by 12 hpl, and a diffuse GFP positive pattern (referred to as matrix), in the outer segment layer and the inner nuclear layer (INL) was present from 15 hpl onwards (Figure 1C), presumably reflecting neutrophil NETosis (Zhu et al., 2021). Analyses of *Tg(mpeg1:mCherry)* and *Tg(mpo:GFP)* double transgenic animals revealed that *mpeg1*:mCherry positive cells co-localized with the GFP positive matrix, suggesting an uptake of the matrix material by monocytes (Figure S1B). T cells of the lymphoid lineage were reported to stimulate Müller glia proliferation, and to augment retina regeneration in a stab wound assay (Hui et al., 2017). To examine if T cells are also recruited to the lesion site, we used transgenic *Tg(lck:NLS-DsRed)* animals, but could not detect any *lck* positive T cells, neither in the homeostatic nor in the regenerating retina up to 7 days post lesion (Figure S2).

**Figure 1:**
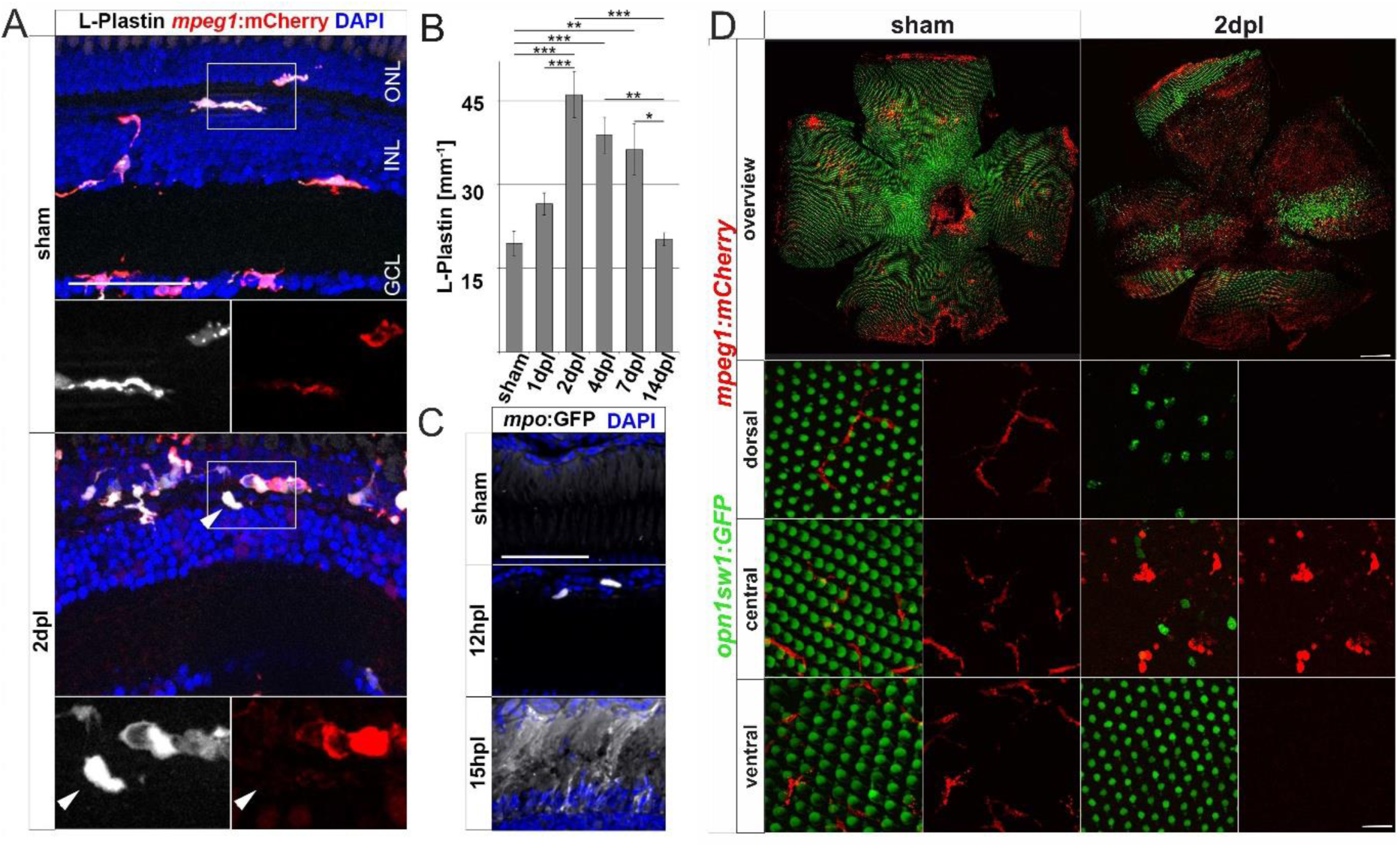
Sterile phototoxic ablation of photoreceptors triggers leukocyte accumulation. (A) Retinal sections show that the majority of L-Plastin^+^ cells are *Tg(mpeg1:mCherry)*^+^, display morphological changes after lesion, and accumulate at the outer nuclear layer (ONL) in response to lesion. However, L-Plastin^+^ *Tg(mpeg1:mCherry)* negative cells were also observed upon injury (arrowhead). (B) Quantification on sections of L-Plastin^+^ cells in sham and regenerating retinae at 1, 2, 4, 7 and 14 days post lesion (dpl) shows an increase of cells within 2dpl that is released within 14 dpl. (C) In contrast to sham, *Tg(mpo:GFP)*^+^ neutrophils are detected upon lesion. They gather at the lesion site (12 hpl) and a diffuse GFP positive pattern (matrix) is formed from 15 hpl onwards. (D) Retinal flat mounts of *Tg(opn1sw1:GFP)* x *Tg(mpeg1:mCherry)* show the ramified structure of leukocytes in a UV-cone-specific reporter line and their reaction to lesion. Upon light lesion, leukocytes increase in number at the lesion site and display an amoeboid, activated morphology and accumulate in the central part of the lesion while unharmed areas appear devoid of *mpeg1:*mCherry positive cells. Scale bars in A & C: 50 µm Scale Bars in D: 200µm, inlets 20µm, error bars indicate standard error; * = p≤0,05; ** = p≤0,01; *** = p<0,001; n≥4; ANOVA Tukey’s post hoc test; INL= inner nuclear layer; GCL=ganglion cell layer.

Taken together, our data show a strong accumulation and activation of innate immune cells following a sterile light lesion that is resolved by 14 dpl, consistent with the mounting of an acute inflammatory response after retinal injury.

### Müller glia activate NF-κB signaling in response to injury

In zebrafish, Müller glia have key functions in the regulation of retinal homeostasis as well as during regeneration (Lenkowski & Raymond, 2014). To study how Müller glia react to injury during regeneration and inflammation, we analyzed the activation of the proinflammatory signaling pathway NF-κB using *Tg(NF-κB:GFP)* and *Tg(gfap:NLS-mCherry)* transgenic animals. The *Tg(NF-κB:GFP)* line reports canonical NF-κB activity via the expression of GFP under the control of six human NF-κB binding motifs driving a c-fos minimal promoter (Kanther et al., 2011). In sham control retinae, *NF-κB*:GFP expression was only found in the retinal vasculature, microglia and, occasionally in Müller glia residing in the inner nuclear layer (Figure 2A). In sharp contrast, robust activation of *NF-κB*:GFP was present in numerous *gfap*:NLS-mCherry positive Müller glia already at 1 dpl (Figure 2A). Moreover, *NF-κB*:GFP expression was present in cells located in the ONL showing the characteristic morphology of photoreceptor cells. Both cell types continued to express *NF-κB*:GFP at 2 dpl. At 4 dpl, *NF-κB*:GFP expression was absent from the ONL but remained detectable in Müller glia. To determine if NF-κB activation might be functionally relevant, we investigated the expression of *matrix metallopeptidase 9* (*mmp9*), a known downstream target of NF-κB with important functions in degrading extracellular matrix and chemokines (Cheng et al., 2012; LeBert et al., 2015; Xu et al., 2018; Yang et al., 2017). Thus, we performed *mmp9 in situ* hybridization in combination with immunohistochemistry against glial fibrillary acidic protein (GFAP/Zrf-1) labeling Müller glia and proliferating cell nuclear antigen (PCNA) labeling cells in S-phase and shortly after. In contrast to sham controls that never showed *mmp9* expression in the homeostatic retina, *mmp9* was strongly expressed by Zrf-1 positive Müller glia at 1 dpl, indicating *de novo mmp9* expression in these cells (Figure 2B). Moreover, numerous *mmp9* and Zrf-1 expressing cells were also positive for PCNA, identifying them as reactively proliferating Müller glia in response to injury. Expression of *mmp9* remained strong at 2 dpl, but dropped below detection levels at 4 dpl, consistent with transcriptome data for *mmp9* (Kramer et al., 2021; Silva 2020; Celotto et al., 2023). Taken together, our results showed that the NF-κB signaling pathway is transiently activated in Müller glia in response to a sterile ablation of photoreceptor cells, indicative of an inflammatory response by Müller glia.

**Figure 2:**
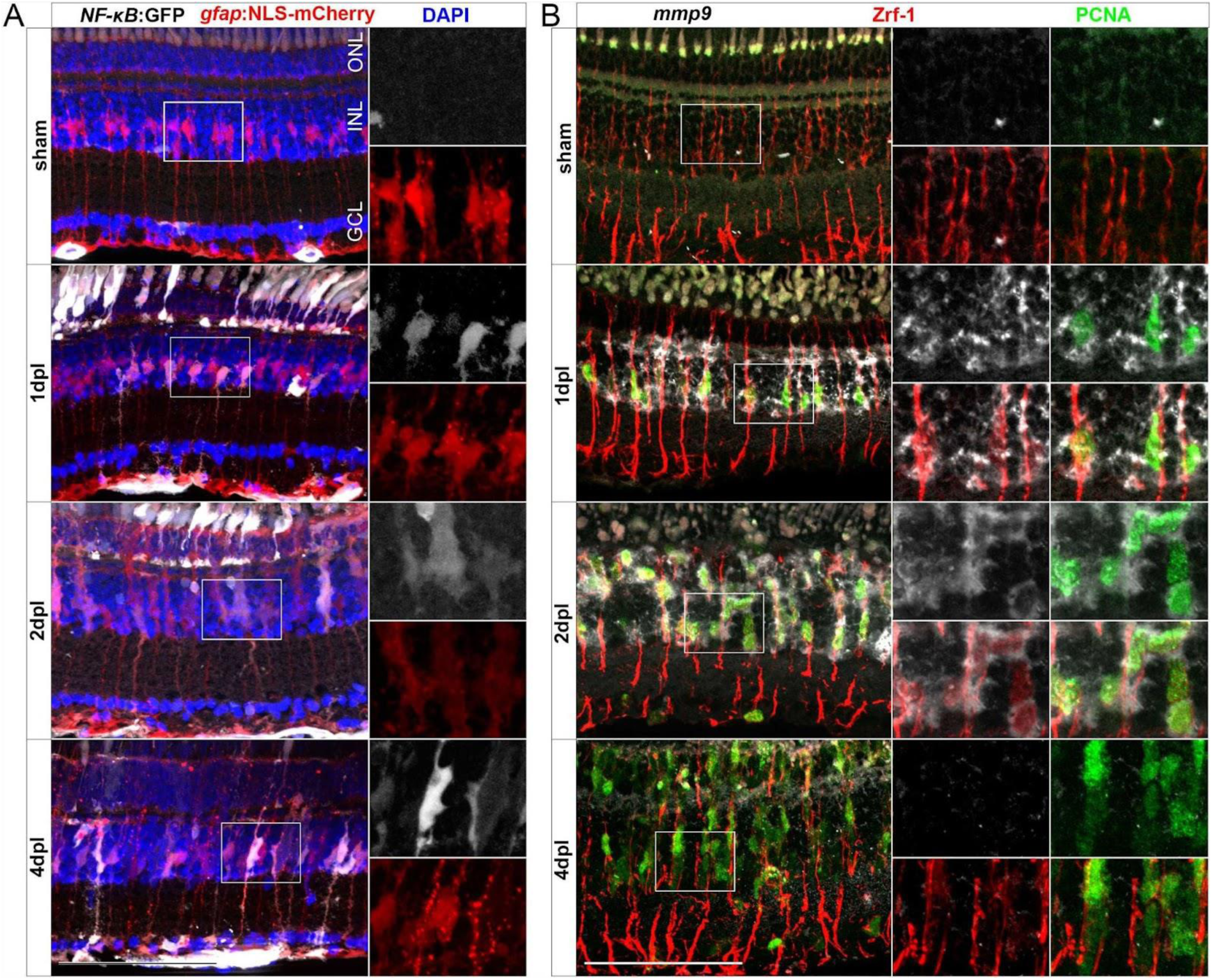
Müller glia transiently activate the *NF-κB:*GFP reporter and express the *NF-κB* target metalloproteinase *mmp9* in response to injury. (A) In comparison to sham, the *NF-κB:*GFP reporter is activated at 1 and 2 days post lesion (dpl) in *gfap*:NLS-mCherry labeled Müller glia localized in the inner nuclear layer (INL) and photoreceptors of the outer nuclear layer (ONL) and is restricted to the INL at 4 dpl. (B) *In situ* hybridization of *mmp9* in combination with immunohistochemistry for proliferating cell nuclear antigen (PCNA) and glial fibrillary acidic protein (GFAP/Zrf-1) labelling Müller glia, shows that *mmp9* is not expressed in sham retinae. In contrast, *mmp9* is transiently expressed at 1 and 2 dpl and returns to undetectable levels at 4 dpl. Scale bar: 50µm. GCL=ganglion cell layer.

### Dexamethasone-mediated immunosuppression reduces retinal regeneration

To study the role of the immune system during retinal regeneration, we used Dexamethasone (Dex), a potent immunosuppressant (Coutinho & Chapman, 2011; Donika Gallina et al., 2015; Kyritsis et al., 2012; Silva et al., 2020; Zhang et al., 2020). Experimental zebrafish were treated with Dex from 10 days prior to lesion until the time point of analysis (Figure 3A). Vehicle control experiments were carried out with the respective amount of the solvent methanol (MeOH). In contrast to MeOH-treated controls, Dex-treatment for 10 consecutive days resulted in an overall reduction of retinal microglia (Figure 3B). Similarly, leukocyte recruitment upon light lesion was significantly reduced in Dex-treated animals at 2 dpl, compared to control (Figure 3B). Quantification of L-Plastin positive cells corroborated a significant reduction at all indicated time points (Figure 3C). In addition, Dex-treatment reduced the number of resident retinal microglia during homeostasis, and no increase in leukocyte number was noted after lesion. To address if the observed changes might be caused indirectly by the known neuroprotective properties of Dex (Gallina et al., 2015), we analyzed cell death using the TdT-mediated dUTP-digoxigenin nick end labeling (TUNEL) assay; however, we did not observe any difference in the number of TUNEL positive cells after light lesion in control MeOH- or Dex-treated animals (Figure S3).

**Figure 3:**
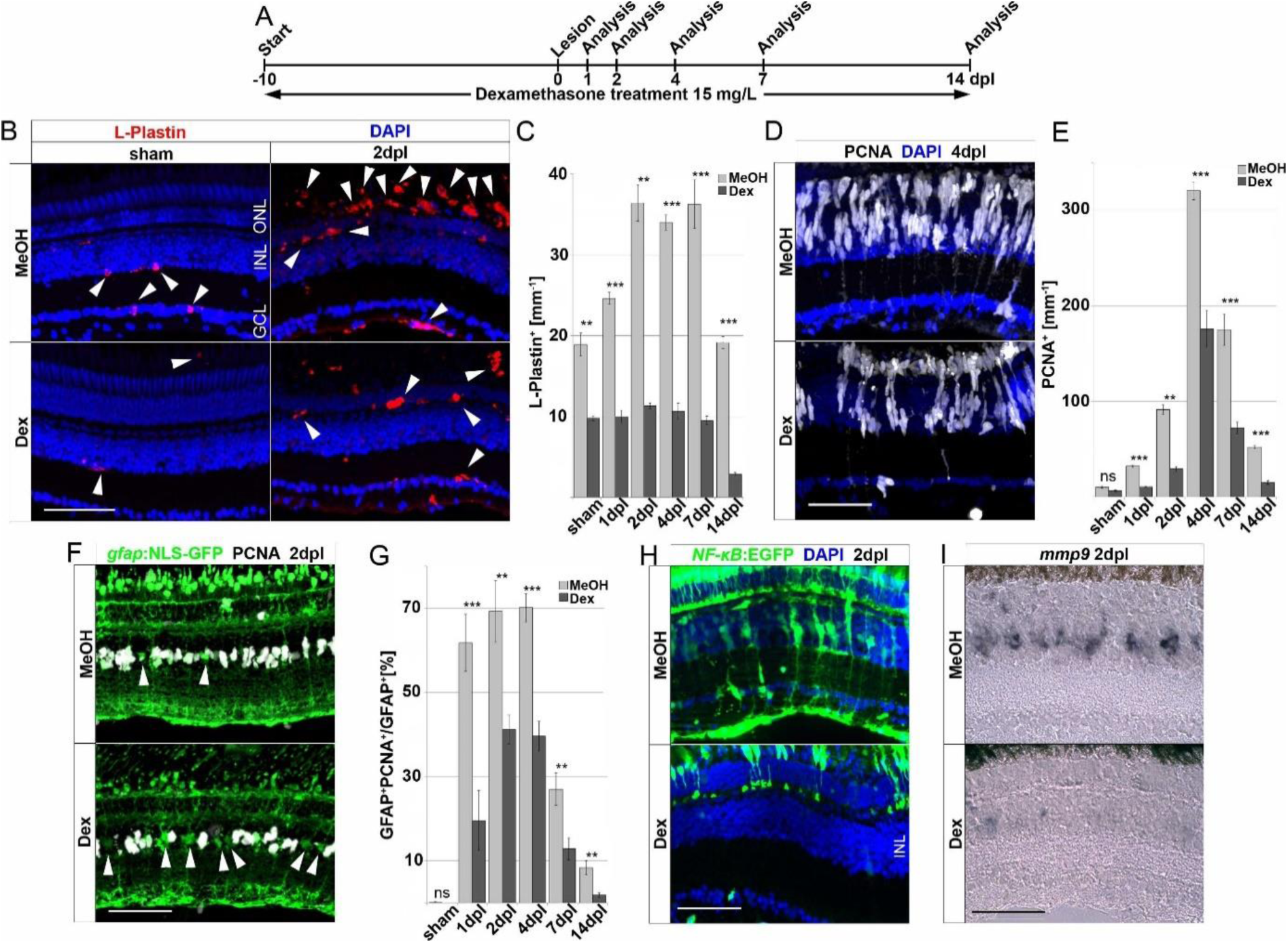
Immunosuppression interferes with leukocyte accumulation and Müller glia reactivity. (A) Scheme of experimental outline. Fish were treated with Dexamethasone (Dex) or vehicle (Methanol; MeOH) from 10 days prior to injury until the day of analysis (sham, 1, 2, 4, 7 and 14 days post lesion; dpl). (B) Dex-treatment reduces the number of L-Plastin^+^ leukocytes (arrowheads) in sham (left panel) and regenerating retinae at 2 dpl (right panel). (C) Quantification of L-Plastin^+^ cells in MeOH- and Dex-treated sham or lesioned animals at 1, 2, 4, 7 and 14 dpl. (D) Immunohistochemistry for proliferating cell nuclear antigen (PCNA) reveals impaired proliferation in Dex-treated animals at 4 dpl. (E) Quantification of PCNA^+^ cells in MeOH- and Dex-treated sham and regenerating retinae at 1, 2, 4, 7 and 14 dpl shows impaired proliferative response in the Dex-treated group. (F) Immunohistochemistry for PCNA in *gfap*:NLS-GFP labeled Müller glia shows reduced numbers of proliferating Müller glia in the Dex treated group at 2 dpl compared to MeOH controls (arrowheads indicating non proliferative Müller cells). (G) Quantification of proliferating Müller glia in vehicle and Dex-treated sham and regenerating retinae at 1, 2, 4, 7 and 14 dpl, indicating that Dex hinders Müller glia proliferation. (H) In comparison to vehicle, the *NFκB*:GFP reporter is not activated in the inner nuclear layer (INL) of Dex-treated retinae at 2 dpl. (I) Injury-induced expression of *mmp9* is strongly reduced in Dex-treated retinae at 2 dpl. Scale bar: 50 µm, Error bars indicate standard error; ns = p>0,05, * = p≤0,05; ** = p≤0,01; *** = p<0,001; n≥6; two-tailed student’s T-Test, GCL=ganglion cell layer, ONL=outer nuclear layer.

Reactive proliferation of Müller glia is a hallmark of retinal regeneration; we therefore further analyzed if Dex-treatment influences proliferation after sterile light lesion using immunolabeling of the proliferation marker PCNA (Figure 3D). In contrast to MeOH-treated controls, Dex-treated animals showed a significant reduction in PCNA positive cells at 4 dpl. Quantification of PCNA positive cells during the course of regeneration revealed that reactive proliferation, driven by Müller cells and neuronal progenitors, is significantly reduced in Dex-treated animals at all time-points examined in comparison to MeOH controls. Furthermore, homeostatic proliferation (most likely by cells of the ciliary margin) in unlesioned retinae is not affected (Figure 3E). To investigate if Dex-treatment specifically affects Müller glia proliferation, we analyzed the number of PCNA positive cells in transgenic *Tg(gfap:NLS-GFP)* animals, which express strong nuclear GFP in all Müller glia (with some leakage of GFP to the cytoplasm, identifying the characteristic Müller glia cell shape, Figure 3F). Indeed, in comparison to MeOH-treated animals, the number of PCNA and *gfap*:NLS-GFP double positive cells was significantly reduced after Dex-treatment. Quantification showed that the percentage of Müller glia co-localizing with PCNA is reduced to less than 50% at all analyzed time points (Figure 3G). This suggests that Müller glia reactivity is impaired upon Dex-treatment. Consistent with this possibility, NF-κB activation is reduced in Müller glia after Dex-treatment, as seen in the *Tg(NF-κB:GFP)* activation reporter line (Figure 3H). As expected, cells in the ONL and Müller glia show a robust *NF-κB*:GFP expression in MeOH-treated control animals at 2 dpl, whereas a reduced number of cells in the ONL activated the *NF-κB*:GFP transgene and no GFP positive Müller glia could be detected at 2 dpl after Dex-treatment. Consistently, activation of the NF-κB downstream target *mmp9* was almost absent in Dex-treated animals at 2 dpl, compared to MeOH controls (Figure 3I). Taken together, these results show that Dex-mediated immunosuppression efficiently reduces the accumulation of leukocytes and impairs Müller glia reactivity, as well as reactive proliferation and regeneration-associated marker gene expression, during retinal regeneration.

### Immunosuppression reduces regeneration of photoreceptors

To examine if the reduced cell proliferation has consequences for the regenerative outcome, we probed the effect of Dex-immunosuppression on restoration of photoreceptors after light lesion. Thus, we performed Dex-treatment and light lesions on *Tg(opn1sw1:GFP)* animals expressing GFP in all UV cones, followed by repeated EdU injections to label newborn cells, and analyzed the animals at 28 dpl when cellular regeneration is completed (Figure 4A). Consistent with the afore-mentioned decrease of retinal leukocytes after Dex-treatment, we observed an almost complete loss of retinal leukocytes after 28 days of continuous immune suppression (Figure S4). In comparison to MeOH-treated control animals, the overall number of EdU positive cells was significantly lower after Dex-treatment at 28 dpl (Figure 4B). Similarly, the number of EdU positive UV-cones (Figure 4C) was severely decreased in Dex-treated animals compared to MeOH-treated controls. Moreover, in addition to a reduction in number, EdU positive UV cones appeared malformed with improperly shaped outer segments in Dex-treated animals (insets Figure 4B). Quantifications showed that the number of EdU positive nuclei was similarly reduced in all retinal layers (Figure 4C). In conclusion, Dex-mediated immunosuppression interferes with reactive proliferation and reactive neurogenesis, indicating that the injury-induced immune response is essential for proper retina regeneration.

**Figure 4:**
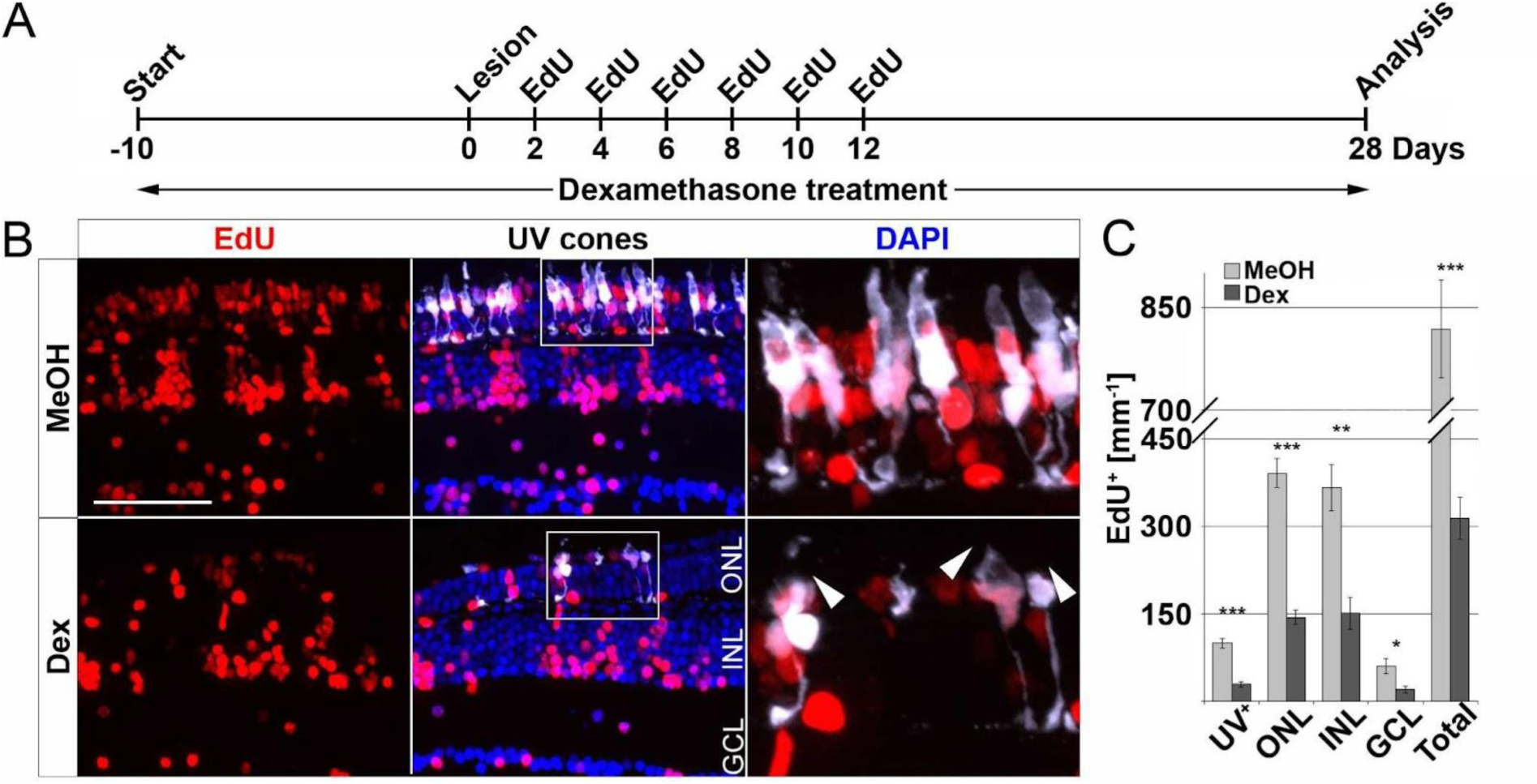
Long-term immune suppression impairs photoreceptor regeneration. (A) Scheme of experimental outline. *opn1sw1*:GFP, labeling UV-cones, transgenic animals were treated with Dexamethasone (Dex) or vehicle (Methanol; MeOH) from 10 days prior to lesion until the day of analysis (28 days post lesion; dpl 38 days post treatment; dpt). EdU pulses were applied at 2, 4, 6, 8 and 10 dpl. (B) In comparison to MeOH-treated animals, the number of regenerated EdU^+^ UV-cones is significantly reduced after Dex-treatment (compare upper panel with lower panel). Furthermore, the morphology of photoreceptor cells is disrupted in comparison to MeOH controls (arrowheads). (C) Quantification of EdU^+^ cells with respect to retinal layers showing significant reduction of EdU positive nuclei in the Dex-treated group. Scale bars: 50 µm. Error bars indicate standard error; * = p≤0,05; ** = p≤0,01; *** = p<0,001; n≥6; two-tailed student’s T-Test. ONL=outer nuclear layer; INL=inner nuclear layer; GCL=ganglion cell layer.

### Retinal microglia support reactive Müller glia proliferation

Activated microglia and macrophages clear debris from dead cells, and they interact with Müller glia and influence their cellular response (Aslanidis et al., 2015; Keightley et al., 2014; Palazzo et al., 2020; Wang et al., 2011). Consistently, our results show that Dex-mediated immunosuppression interferes with reactive proliferation and neurogenesis, indicating that the injury-induced immune response is essential for successful retina regeneration. To independently test this notion, we investigated whether microglia positively contribute to reactive proliferation during retina regeneration, using Interferon regulatory factor 8 (*irf8*) myeloid-defective mutants to genetically deplete microglia in embryonic and juvenile fish (Shiau et al., 2015). To verify leukocyte deficiency at adult stages, we analyzed L-Plastin immunoreactivity in homozygous adult *irf8* mutants and heterozygous control siblings, and found that the number of leukocytes in the homeostatic retina was significantly reduced in *irf8* mutants (Figure 5A, B). Next, we asked whether the microglia deficiency also affects the proliferation of Müller cells and NPCs during regeneration. We therefore combined the light lesion paradigm with EdU-labeling at 3 dpl prior to analysis at 4 dpl (Figure 5C). Compared to heterozygous control siblings, the number of accumulated L-Plastin positive cells was clearly reduced in *irf8* mutant animals at 4 dpl (Figure 5D). Similarly, the number of EdU positive cells is lower in *irf8* mutants at 4 dpl compared to heterozygous controls (Figure 5D). Quantification of both the number of L-Plastin positive cells as well as the number of EdU positive cells confirmed a statistically significant reduction (Figure 5E). Thus, consistent with our above findings using Dex-mediated immunosuppression, these results show that genetic depletion of leukocytes correlates with decreased reactive proliferation upon injury, supporting the notion of an important role of immune cells during zebrafish retina regeneration.

**Figure 5:**
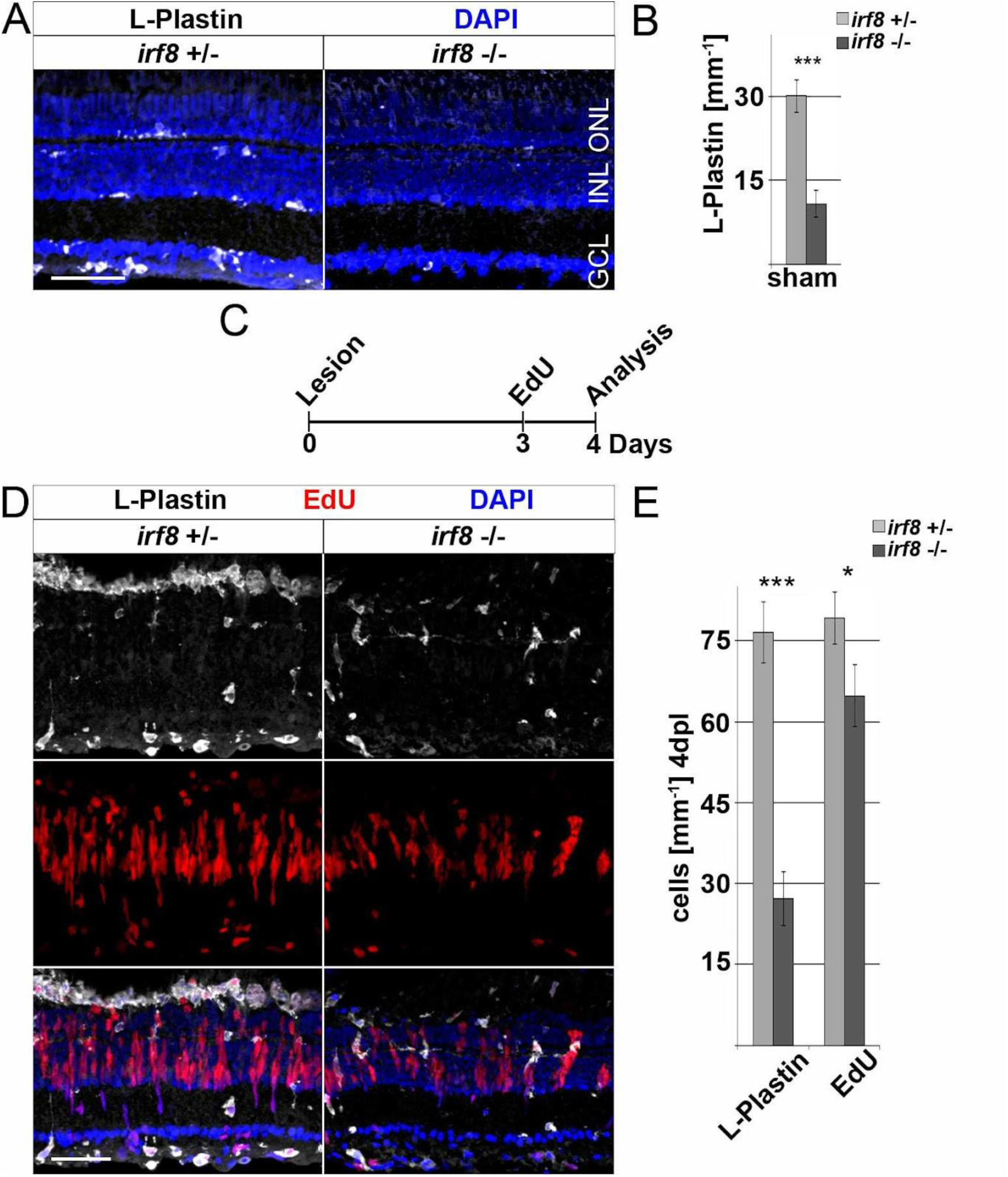
Genetic reduction of microglia inhibits retinal proliferation in response to injury. (A) In contrast to heterozygous control siblings (*irf8*^+/-^), interferon regulatory factor 8 (*irf8)* homozygous myeloid-deficient mutant retinae (*irf8*^-/-^) display a decreased number of L-Plastin^+^ leukocytes in homeostasis. (B) Quantifications of L-Plastin^+^ cells in *irf8*^-/-^ and control heterozygous siblings *irf8*^+/-^. (C) Scheme of experimental outline. *irf8*^-/-^ and *irf8*^+/-^ animals received an EdU pulse at 3 days post lesion (dpl) and were analyzed at 4 dpl. (D) During regeneration, the accumulation of L-Plastin^+^ cells at the outer nuclear layer (ONL) is impaired in *irf8^-/-^* but not in control *irf8^+/-^* siblings. Moreover, the number of EdU^+^ cells is significantly decreased in *irf8*^-/-^ retinae. (E) Quantification of L-Plastin^+^ cells and EdU^+^ nuclei in *irf8*^-/-^ and control *irf8*^+/-^ animals at 4 dpl, show reduced amounts of positive cells respectively. Scale bar: 50 µm. Error bars indicate standard error; * = p≤0,05; *** = p<0,001; n=5; two-tailed student’s T-Test; INL=inner nuclear layer; GCL ganglion cell layer.

### Inflammatory stimuli trigger a regeneration response in Müller glia in the absence of lesion

We hypothesized that inflammation might not only be necessary, but also sufficient, to trigger reactive proliferation of Müller glia and the generation of NPCs. To address this notion, we injected different immune activators into the vitreous of the eye - thus leaving the retina unlesioned (Fig 6A) - and analyzed the response of Müller glia at 2 and 4 days post injections (dpi; Figure 6). The toll-like receptor (TLR) agonist zymosan has previously been shown to stimulate an inflammatory response both in the zebrafish and mouse eye, and in the zebrafish telencephalon (Kyritsis et al., 2012; Kurimoto et al., 2013; Zhang et al., 2020). Next, we tested injections of flagellin, the principal structural protein of bacterial flagella, which is known to cause sepsis in zebrafish larvae (Barber et al., 2016). As a control, to ensure that injection of both zymosan and flagellin did not cause retinal cell death and thereby trigger a regenerative response indirectly, we performed TUNEL assays at 2 dpi. Neither zymosan nor flagellin injection resulted in a significant increase of TUNEL positive cells in comparison to control PBS-injected eyes (Figure 6B). Interestingly, the number of L-Plastin positive cells also did not significantly increase after injecting zymosan or flagellin (Figure 6C). In contrast, cell proliferation using PCNA immunohistochemistry is clearly induced in both zymosan as well as flagellin-injected animals in comparison to PBS-injected controls (Figure 6D). Quantification of PCNA positive nuclei confirmed a significant increase in zymosan and flagellin-injected animals at 4 dpi, but not at 2 dpi (Figure 6E). Furthermore, we analyzed activity of NF-κB after injections of PBS, zymosan and flagellin using the *NF-κB*:GFP reporter line. Consistent with the afore-mentioned expression in non-injected retinae, PBS-injected controls show *NF-κB*:GFP expression in blood vessels, microglia and few Müller glia residing in the INL (Figure 6F). In contrast, *NF-κB*:GFP is strongly activated in additional cells of the inner and outer nuclear layer in zymosan and flagellin-injected animals at 2 dpi. Again, GFP positive cells in the ONL showed mostly the characteristic morphology of photoreceptor cells (see also above, Fig. 2B & 3H), in particular in the zymosan-injected animals. Consistent with the activation of the *NF-κB*:GFP reporter, retinal expression of *mmp9* is also found in zymosan and flagellin-injected animals, but not in PBS-injected controls at 2 dpi (Figure 6G). Finally, we analyzed the expression of the transcription factor *her4.1*, a downstream target of the Notch signaling pathway, which is upregulated in NPCs generated in response to photoreceptor damage (Wan et al., 2012). No expression of *her4.1* is detected in PBS-injected controls, but *her4.1* expression is activated in zymosan and flagellin-injected samples at 4 dpi (Figure 6H). These results show that activators of the immune system can strongly stimulate Müller glia reactivity and proliferation, even in the absence of tissue damage.

**Figure 6:**
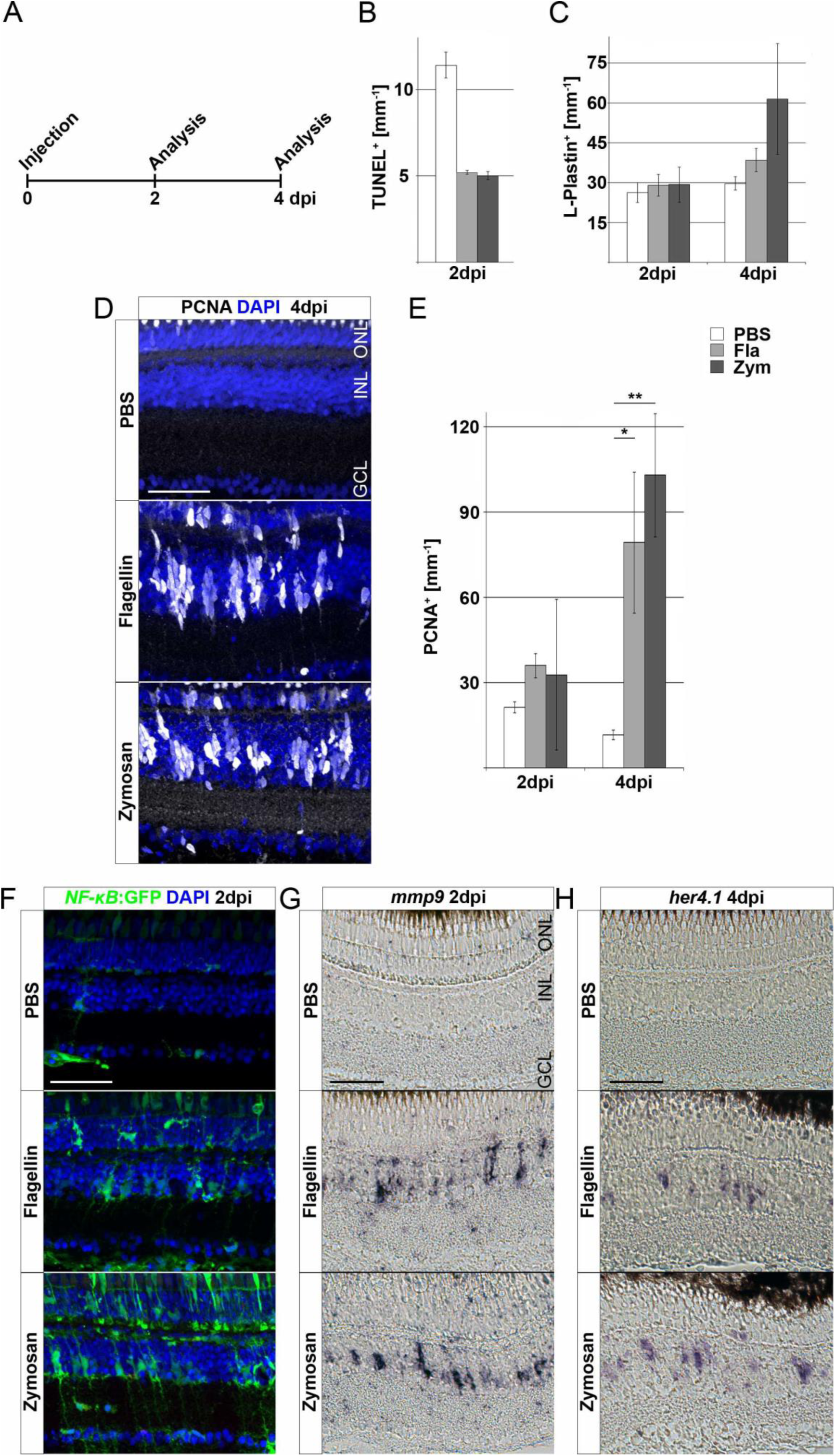
Inflammatory stimuli trigger Müller glia reactivity. (A) Scheme of experimental outline. Wild type or *NF-κB*:GFP reporter animals were injected with flagellin or zymosan into the vitreous of the eye and analyzed at 2 and 4 days post injection (dpi). (B) Quantification of TUNEL^+^ nuclei in control and injected retinae at 2 dpi revealing no increase in cell death. (C) Quantification of L-Plastin^+^ cells in control (PBS) and flagellin or zymosan injected retinae at 2 and 4 dpi display no significant increase in leukocytes due to the injection of factors. (D) Immunohistochemistry for proliferating cell nuclear antigen (PCNA) reveals proliferation in flagellin or zymosan injected but not in control animals at 4 dpi. (E) Quantification of PCNA^+^ nuclei in injected retinae at 2 and 4 dpi. (F) In comparison to PBS injected animals, injection of flagellin or zymosan results in strong activation of the *NFκB*:GFP reporter at 2 dpi. (G & H) *In situ* hybridization of *mmp9* and *her4.1* show expression of both genes in flagellin or zymosan injected but not in PBS injected eyes at 2 and 4 dpi, respectively. Scale bar: 50 µm. Error bars indicate standard error; * = p≤0,05; ** = p≤0,01; *** = p<0,001; n=6; two-tailed student’s T-Test; ONL=outer nuclear layer; INL=inner nuclear layer; GCL=ganglion cell layer.

### M-CSF injection triggers leukocyte accumulation and Müller glia reactivity in the absence of a lesion

In order to identify the stimulatory potential of specific individual inflammatory mediators, we focused on macrophage colony-stimulating-factor (M-CSF/CSF-1), based on preliminary data from our recent single-cell RNAseq dataset of regenerating retina (Celotto et al., 2023), and using the same injection paradigm as before (Figure 7A). In mammals, M-CSF stimulate CSF-1 receptor signaling, which is involved in monocyte colonization and stimulation (Chitu et al., 2016; Wu et al., 2018). Injection of human M-CSF induced an increased cell proliferation at 4 dpi compared to PBS controls (Figure 7B, C), despite the absence of unspecific damage to the retina, as seen by lack of an increase in TUNEL positive cells at 2 dpi. Quantification corroborated a significant increase of proliferating cells, scored as PCNA positive nuclei, in M-CSF-injected retinae, compared to PBS-injected controls (Figure 7D). To address if M-CSF causes an increase in the number of retinal leukocytes, we performed immunolabeling of L-Plastin in M-CSF- and PBS-injected specimens. We observed a significant accumulation of L-Plastin positive cells in M-CSF-, but not in PBS-injected, animals at 4 dpi (Figure 7E & F). Further, we found augmented NF-κB activity after M-CSF injection in *NF-κB*:GFP reporter animals. PBS-injected controls showed the homeostatic *NF-κB*:GFP expression described above (Figure 7G). In sharp contrast, additional *NF-κB*:GFP positive cells were found in the M-CSF-injected animals at 2 dpi, located in the inner nuclear layer and showing the characteristic morphology of Müller glia. Photoreceptor cells in the ONL did not activate the *NF-κB*:GFP reporter following M-CSF injection whereas they do so upon lesion (see above, Fig. 2B & 3H), which we tentatively suggest to reflect a more selective role of M-CSF signaling for Müller glia. Consistent with the activation of *NF-κB*:GFP, *mmp9* was also found to be expressed in M-CSF injected animals, but not in PBS-injected controls at 2 dpi (Figure 7H). Taken together, these data demonstrate that human recombinant M-CSF can induce leukocyte accumulation in the neuronal retina of zebrafish, and can stimulate Müller glia reactive proliferation and alter gene expression even in the absence of a lesion.

**Figure 7:**
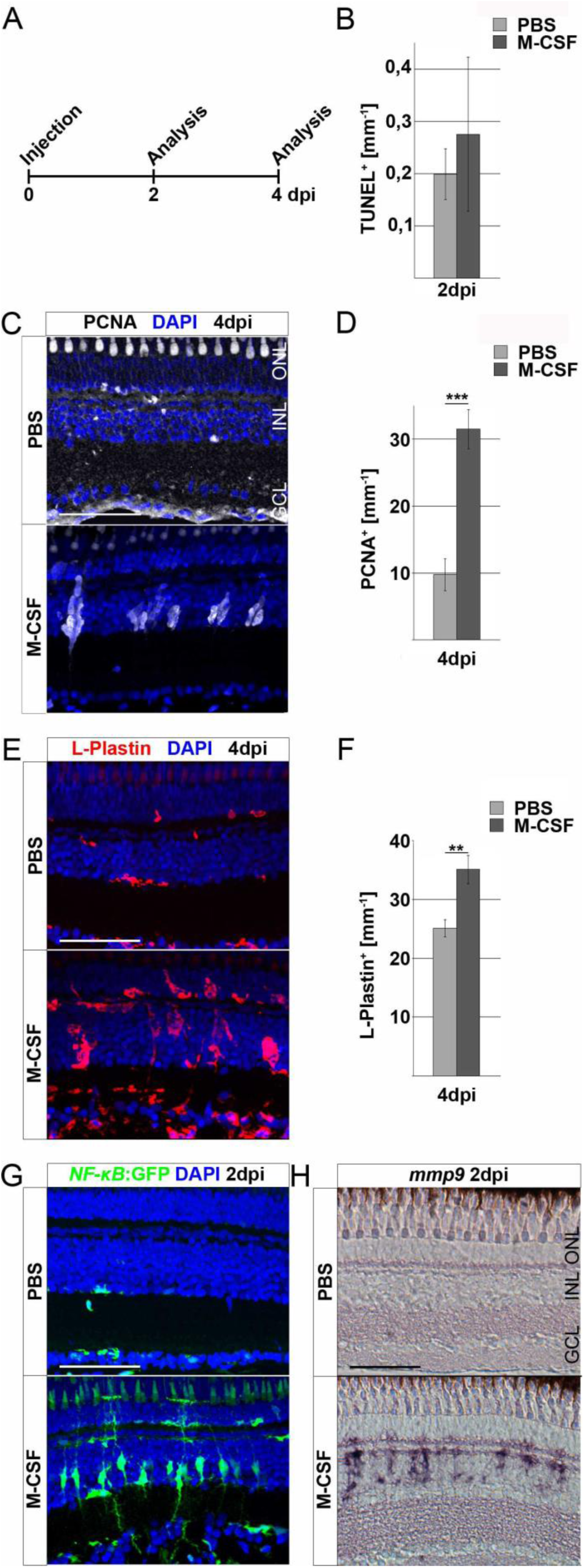
M-CSF stimulates an inflammatory response and initiates Müller glia cell cycle re-entry in the absence of lesion. (A) Scheme of experimental outline. Wild type or *NFκB*:GFP reporter animals were injected with M-CSF or PBS into the vitreous of the eye and analyzed at 2 and 4 days post injection (dpi). (B) Quantification of TUNEL^+^ nuclei in control and M-CSF-injected retinae at 2 dpi reveals no chance in cell death. (C) Immunohistochemistry for proliferating cell nuclear antigen (PCNA) reveals proliferation in M-CSF injected but not in control injected animals at 4 dpi. (D) Quantification of PCNA^+^ cells in control (PBS) and M-CSF injected retinae at 4 dpi. (E) L-Plastin staining shows more positive cells in M-CSF injected retinae at 4 dpi compared to controls. (F) Quantification of L-Plastin^+^ cells in control and M-CSF injected retinae at 4 dpi, show accumulation of L-Plastin cells in M-CSF injected specimen. (G) In comparison to PBS, injection of M-CSF results in strong activation of the *NFκB:*GFP reporter at 2 dpi. (H) *In situ* hybridization of *mmp9* shows transcriptional activation of this gene in M-CSF injected but not in control-injected animals at 2 dpi. Scale bars = 50µm, Error bars indicate standard error; ** = p≤0,01; *** = p<0,001; n>5; two-tailed student’s T-Test. ONL=outer nuclear layer; INL=inner nuclear layer; GCL=ganglion cell layer.

## Discussion

In contrast to mammals, the zebrafish retina readily regenerates photoreceptor cells that are lost following a phototoxic lesion. Several signaling pathways have been implicated in the restoration of lost neurons (Lahne et al., 2020; Lenkowski & Raymond, 2014; Goldman, 2014; Hochmann et al., 2012). A role for the immune system in regeneration of the adult zebrafish retina has previously been suggested (Mitchell et al., 2018; Sifuentes et al., 2016; Silva et al., 2020; White et al., 2017; Zhang et al., 2020; Iribarne et al., 2022), but it is not understood in detail. Here, we confirm and extend these previous studies on the role of inflammation, and its impact on Müller glia reactivity, by modulating immune activity during regeneration of the adult zebrafish retina after a sterile phototoxic ablation of photoreceptor cells. We find that microglia and neutrophils infiltrate and accumulate at damaged sites of the retina. Importantly, Müller glia themselves react by activating the proinflammatory NF-κB signaling pathway in response to retinal damage. Our functional studies show that Dexamethasone-mediated immunosuppression reduces (i) leukocyte accumulation, and (ii) the reactive Müller glia response at the proliferative and gene expression level, and thus reduces the regeneration of photoreceptors. (iii) Conversely, in gain-of-function assays, injection of flagellin, zymosan or M-CSF as inflammatory factors that are thought to facilitate the immune system, some Müller glia are stimulated and reactive proliferation is induced (Gorsuch & Hyde, 2014; Lenkowski et al., 2013, Nelson et al., 2013; Figure 8). Overall, we suggest that cells of the innate immune system act as necessary and sufficient positive regulators for successful regeneration of the adult zebrafish retina. Regulators of the inflammatory state, such as Dexamethasone and the M-CSF that stimulates Müller glia-based photoreceptor regeneration in our injection assays, might therefore provide advanced options for clinical treatments of retinal disease, as recently also suggested for corticosteroid management of early phases of spinal cord injury (Nelson et al., 2019).

**Figure 8:**
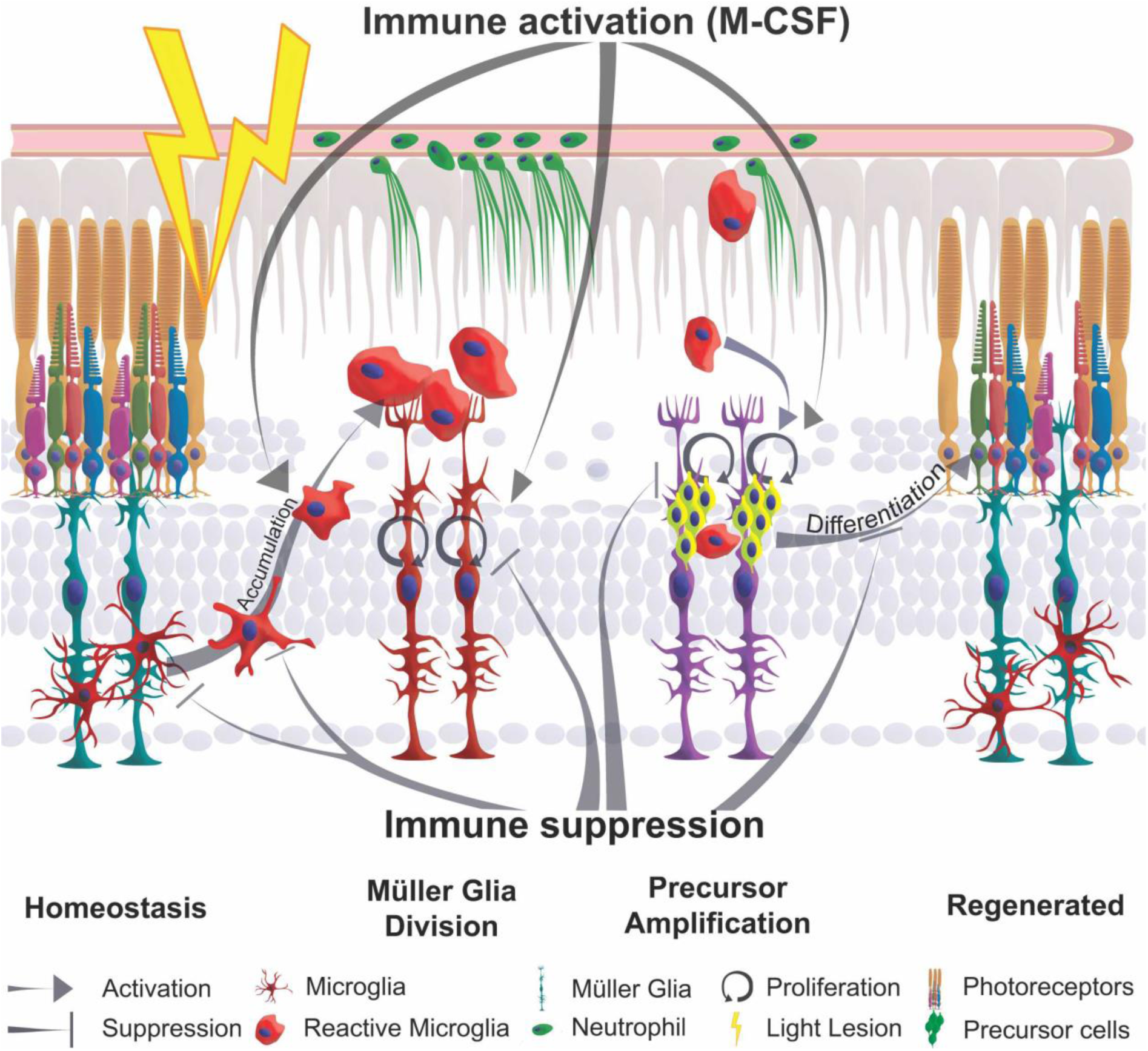
Schematic summary of the impact of inflammation in retinal regeneration. In response to injury, microglia undergo a phenotypic change from amoeboid to ramified shape and accumulate at the lesion site. This process is inhibited by immune suppression. The reactive proliferation of Müller glia during regeneration is also impaired upon immunosuppression. Leukocytes stimulate the proliferation of neuronal precursor cells. M-CSF as inflammatory mediator triggers Müller glia proliferation, leukocyte accumulation and the generation of neuronal precursor cells. This process may also act indirectly via the stimulation of immune cells. Overall, the innate immune system, via mediators such as M-CSF, supports the differentiation into photoreceptor cells and thereby acts as a regenerative cue for retina regeneration.

### Sterile ablation of photoreceptors recruits innate immune cells

Consistent with previous studies, we find that retinal *mpeg1*:mCherry positive tissue-resident microglia display a ramified morphology in homeostasis, but change to an activated amoeboid cell shape upon a sterile light lesion, similar to what is observed after CNS lesion (Karlstetter et al., 2010; Kroehne et al., 2011; Kyritsis et al., 2012; Mitchell et al., 2018). Along with activation, we find a strong accumulation of microglia at the lesion site, which is resolved over a period of 14 days. The underlying dynamic events indicate an acute inflammatory response of leukocytes that is tightly controlled in a spatiotemporal manner. We assume that it is mostly tissue-resident microglia that mediate the observed immune response; however, we cannot exclude that additional peripheral macrophages are also recruited from the bloodstream and contribute to the observed reaction, in particular since we also observe neutrophils entering the retina. Consistent with our assumption, live imaging in larval zebrafish revealed that peripheral macrophages do not enter the developing retina during drug-mediated ablation of rod photoreceptors (Oosterhof et al., 2017; White et al., 2017). In contrast, neurotoxic ablation of the ganglion cells and neurons of the inner nuclear layer using ouabain triggers proliferation of adult retinal microglia, and it has been suggested that the observed increase in cell number might also involve peripheral macrophages entering the retina from circulation (Mitchell et al., 2018). The observation of *mpo*:GFP positive neutrophils in the vasculature, as well as neutrophil accumulation in the photoreceptor outer segment layer upon light lesion, indicates a certain permeabilization of the blood-retina-barrier, which may also allow additional leukocytes to invade the damaged retina (Eshaq et al., 2017; McMenamin et al., 2019; Sim et al., 2015). Further studies of the cell-permeability of the blood-retina-barrier during regeneration, e.g. by *in vivo* imaging approaches, and specific labeling of either peripheral macrophage or microglia populations, will likely be interesting in this regard.

Our observation that neutrophils arrive in the retina following non-invasive light lesion suggests an interesting aspect of retina regeneration, with parallels to observations after neurotoxic ablation of retinal ganglion cells (Mitchell et al., 2018). Neutrophils have different functions at the site of injury, and they are among the first cells to react to injury. It has been proposed that they participate in clearing cellular debris, by secretion of factors influencing angiogenesis and regeneration, but they may also support resolution of inflammation (Wang, 2018). In mice, neutrophils promote the regeneration of the optic nerve by expression of oncomodulin, and can influence the response of glial cells and axonal outgrowth after spinal cord injury (Kurimoto et al., 2013; Stirling et al., 2009; Tsarouchas et al., 2018). Furthermore, neutrophils are indicated to resolve inflammation by a localized O_2_ depletion (Campbell et al., 2014). Thus, neutrophils could be involved in the mediation of the immune response in adult retina regeneration in zebrafish upon lesion. Further investigation of the observed GFP positive matrices may therefore be interesting. Here, we speculate that these structures might act as ‘extracellular-traps’ that could prevent cellular debris from spreading further into the surrounding tissues, protecting the latter from secondary cell death (Branzk & Papayannopoulos, 2013; Walker et al., 2007), which might help suppress a systemic inflammatory/regenerative reaction in the retina. The observation of GFP positive inclusions in mpeg1 positive macrophages furthermore suggests degradation of the trap-like structures by macrophage uptake (Farrera & Fadeel, 2013).

While microglia and neutrophils seem to play important roles in retina regeneration, we did not observe any infiltration of *lck*:NLS-dsRed positive T cells into the retina after lesion, arguing that T cells may be dispensable at least for the initial regeneration response. A recent report suggests that T cells can positively influence the regenerative response after an invasive mechanical lesion of the retina (Hui et al., 2017); this difference might be due to differences in the lesion paradigm employed, and/or timing of the regeneration events. As mentioned above, the non-invasive light lesion that we employed results in a specific ablation of photoreceptors without physically disrupting the blood-retina barrier. In contrast, the needle stab employed by Hui et al., 2017 disrupts the blood-retina barrier, thereby allowing T cells to access the lesion.

### Müller glia react to photoreceptor cell death in an inflammatory manner

Upon retinal injury and progressive dystrophies, mammalian Müller glia undergo physiological changes and show transcriptional alterations as a hallmark of their gliotic response (Bringmann et al., 2009; Bringmann & Wiedemann, 2011). Despite the regenerative response of zebrafish Müller glia, they exhibit initial signs of reactive gliosis following retinal damage (Thomas et al., 2016). Here, we report that ablation of photoreceptors induces an acute inflammatory response at the molecular level. Importantly, we observed an inflammatory-like response directly in the lesion-responding Müller glia. Using *NF-κB*:GFP reporter fish (Kanther et al., 2011), we find that canonical NF-κB becomes activated in Müller glia upon damage, in keeping with transcriptomic changes of Müller glia upon light lesion (Sifuentes et al., 2016; Celotto et al., 2023). In the regenerating chick retina, NF-κB is active in Müller glia and acts as an important pathway controlled by factors that are secreted by microglia (Palazzo et al., 2020). A potential activator of NF-κB may be TNFα; indeed, several studies suggested that dying neurons secrete this factor and thus trigger Müller glia proliferation and the generation of NPCs (Kumar & Shamsuddin, 2012; Nelson et al., 2013).

Consistent with previous reports, we also found the NF-κB target gene, matrix metallopeptidase *mmp9,* to be expressed in reactive Müller glia, displaying a highly dynamic and strictly spatiotemporally controlled activation profile (Kaur et al., 2018; Kim et al., 2007; Silva et al., 2020; Takada et al., 2004). In *mmp9* knockout zebrafish, Müller glia-derived progenitors show a hyperproliferative response after injury, but reduced survival of regenerated photoreceptors (Silva et al., 2020). The exact function of *mmp9* is not completely clear yet; we speculate that *mmp9,* as a metalloproteinase, might be involved in the degradation of extracellular matrix (ECM), thus ‘opening’ the retina for incoming immune or neuronal precursor cells, and/or modulating ECM interacting cytokines (Gong et al., 2008; LeBert et al., 2015; Xu et al., 2018; Yoong et al., 2007). Interestingly *mmp9* is a key marker gene known to be differentially expressed between wet and dry forms of age-related macular degeneration, providing further evidence for a possibly critical role during inflammatory processes (Fritsche et al., 2016). Understanding the contribution and molecular control of *mmp9* in zebrafish retina regeneration may therefore provide further insights into the mechanisms and molecules that are required to restore lost neural tissues.

### Immunosuppression alters the regenerative response of the zebrafish retina

Immunosuppression with the glucocorticoid dexamethasone drastically decreases the amount of retinal microglia and blocks accumulation of reactive leukocytes in the retina after lesion (Gallina et al., 2014; White et al., 2017). In addition, activation of NF-κB signaling was also successfully blocked in Müller glia, as shown by the loss of *NF-κB*:GFP and expression of its target gene. NF-κB activation has been described in a plethora of cellular functions such as cell survival, cell death and mediation of inflammation by the regulation of cytokine expression, as well as cell proliferation and migration (Liu et al., 2017). In Müller glia, NF-κB signaling could mediate an inflammatory-like response, consistent with their expression of pro- and anti-inflammatory cytokines (Nelson et al., 2013; Polager & Ginsberg, 2002; Zhao et al., 2014). Consistently, following Dex-treatment, we find that the reactive proliferation of Müller glia was strongly reduced, indicating that inflammatory-like signaling promotes Müller glia cell cycle reentry. Interestingly, homeostatic proliferation, which mainly occurs in the ciliary marginal zone and only sporadically in the central retina, remains unaffected, similar to our findings in the zebrafish telencephalon (Kyritsis et al., 2012). Moreover, Dex-treatment is not only affecting reactive proliferation, as is also confirmed by other research groups (Iribarne & Hyde, 2022; Silva et al., 2020; White et al., 2017). Regenerated UV-cones show dysmorphic outer segments, indicating that Dex-treatment also impairs the proper differentiation of photoreceptor cells. On the other hand, in mmp9 mutants, initial Dex-treatment supports the restoration of the photoreceptors and thus can act positively (Silva et al., 2020). Collectively, we propose that inflammatory signaling is required for cell cycle reentry of Müller glia during regeneration, but might also play additional roles during maturation of regenerated photoreceptors (Figure 8). An incomplete regenerative outcome was also detected in larval retinal pigment epithelium (RPE) regeneration in *irf8*-mutants deficient in microglia. Upon ablation of RPE, the neural retina showed an increased reactive proliferation in mutants, but the amount of regenerated RPE cells is lower than in wild type siblings (Hanovice et al., 2019).

To further understand the impact of resident microglia on the regeneration process in adult retina, we addressed the proliferative potential of retinae in response to injury in *irf8* mutant fish, as a genetic model for myeloid deficiency in larval and juvenile zebrafish (Shiau et al., 2015). We confirm microglia deficiency at adult stages (>6 month), indicating that *irf8* is required throughout lifetime in primitive as well as a definitive wave of monocyte populations of the eye (Ferrero et al., 2018; Shiau et al., 2015). However, a reduced number of leukocytes is still found to populate the adult retina. Upon lesion, we observed that the number of cells that are in cell cycle is significantly reduced in *irf8* mutants in comparison to heterozygous siblings in response to lesion. Interestingly, the number of L-Plastin+ leukocytes in *irf8* mutants increases upon injury, but never reaches levels comparable to heterozygous control siblings. In line with other reports, these findings indicate that microglia are supportive for Müller cell proliferation and NPC amplification and thereby support initial steps of regeneration (Conedera et al., 2019; Zhang et al., 2020; Iribarne & Hyde, 2022). The necessity of (tissue-resident) macrophages is also observed in other tissues, e.g. telencephalon, spinal cord, caudal fin and heart, showing the contribution of the innate immune system to tissue regeneration (Kyritsis et al., 2012; De Preux Charles et al., 2016; Petrie et al., 2015; Portillo et al., 2017; Roche et al., 2018; Tsarouchas et al., 2018). This supportive role of macrophages in zebrafish is unlike that of the rodent retina, where microglia appear to inhibit Müller-glia-mediated tissue regeneration by suppression of *ascl1* expression. This interesting species difference might be explained by distinct expression profiles of zebrafish vs. mouse microglia during the time course of regeneration (Issaka Salia & Mitchell, 2020; Mitchell et al., 2019; Todd et al., 2020).

### Immune stimulation triggers Müller glia reactivity and proliferation

Retinal pathology is often coupled to neuronal loss, activation of the immune system and Müller glia reactivity (Iribarne & Hyde, 2022). In particular, mammalian Müller glia react to injury mostly by (proliferative) gliosis, which contributes to glial scarring, thus enhancing loss of vision (Bringmann et al., 2006). In zebrafish, retinal injury also results in an acute immune reaction; however, in contrast to mammals, this is followed by a regenerative response.

Importantly, inflammatory signals can indeed initiate reactive proliferation, even in the absence of injury, as seen by our vitreous injections of toll-like receptor (TLR) agonists zymosan and flagellin to trigger an inflammatory response (Akira & Takeda, 2004; Barber et al., 2016; Zhang et al., 2020). Leukocyte number was not increased following these injections, which might be explained by three possible scenarios: i) Zymosan and flagellin do not stimulate leukocyte proliferation ii) recruitment from the bloodstream is not induced; while in both scenarios a local leukocyte activation can occur. iii) The accumulation could already be resolved within 48h, since the response of immune cells is rapid and highly dynamic. Furthermore, the injected concentration of zymosan in a rodent model, where leukocytes are seen to accumulate, is ten-fold higher than the concentration used in our injections, and could reflect a dose dependency (Kurimoto et al., 2013). In contrast to the more generic inflammatory stimuli, M-CSF-injected retinae show an increased number of leukocytes. This may be the consequence of both proliferation and recruitment from the bloodstream, since M-CSF receptor (CSF-1R) is thought to regulate the population of microglia in neural tissue during development (Wu et al., 2018). Following M-CSF injection, we find that Müller glia activate NF-κB signaling, which fits well with studies reporting TLR and CSF-1 receptor expression in Müller glia and microglia (Bsibsi et al., 2002; Kumar & Shamsuddin, 2012; Olson & Miller, 2004; Oosterhof et al., 2017; Rocío Nieto-Arellano, 2019). Most importantly, we found that in the absence of injury, stimulation of the immune system activates the regenerative response program of Müller glia, as shown by upregulation of *mmp9* and *her4.1,* as well as an increase in proliferation; consistently, expression of *her4.1* also indicates the generation of NPCs from Müller glia. The expression of the CSF-1 receptor in neural progenitor cells is also associated with proliferation, survival and differentiation, supporting the theory that M-CSF stimulates a proliferative/regenerative response in Müller glia (Chitu et al., 2016). Nevertheless, whether Müller glia reactivity is a direct consequence of CSF/TLR-agonist treatment, or an indirect consequence of potentially altered microglia signaling, is so far not clear, because receptors for CSF/TLR signaling appear to be expressed in both cell populations (Kochan et al., 2012; Kumar & Shamsuddin, 2012; Letiembre et al., 2007; Lin et al., 2013; Mitchell et al., 2019; Tsarouchas et al., 2018; Wu et al., 2018). Hence, it will be interesting to test whether CSF-1R or TLR agonists alone are sufficient to drive increased Müller glia proliferation, using a model that completely lacks microglial cells in the retina. Moreover, it will be interesting to determine if the observed effects of the CSF1R/TLR-agonists are mediated via NF-κB signaling, or via other pathways, such as the MAP kinase pathway, or combinations thereof (Chen et al., 2018; Kawasaki & Kawai, 2014; Wan et al., 2012).

## Conclusion

Our work strongly supports the notion that acute inflammatory signaling provides a beneficial contribution during adult zebrafish retina regeneration, akin to findings in the telencephalon (Kyritsis et al., 2012; Kizil et al., 2015). Reactivity of retinal leukocytes appears to be tightly controlled after injury, and the presence of leukocytes positively influences reactive proliferation of Müller glial cells, as well as the downstream differentiation of Müller glia progeny into mature photoreceptor cells. Importantly, immune stimulation through TLRs, or via M-CSF injection, was sufficient, in the absence of injury, to trigger Müller glia proliferation and NPC formation. Hence, we conclude that inflammatory signaling plays a critical role during zebrafish retina regeneration.

Further investigations on possible interactions between leukocytes and Müller glia will help to understand the balance between the inflammatory reaction and its resolution during regeneration of neural tissue in zebrafish. In addition, it will be important to identify critical factors secreted by leukocytes in an acute reaction to lesion that are - directly or indirectly-involved in triggering Müller glia reactivity, and thus, regeneration. Finally, this study provides a framework for investigating potentially beneficial effects of an acute immune response, also in comparison with failed regeneration and chronic inflammatory outcome in mammalian models.

## Materials and Methods

### Ethics statement

The animal experiments were performed in strict accordance with the European Union and German law (Tierschutzgesetz). Experimental procedures were approved by the animal ethics committee of the TU Dresden and the Landesdirektion Sachsen (permits AZ: 24-9168.11-1/2013-5; AZ: TV A 1/2017; TVV 21/2018; AZ: TVV 55/2018).

### Zebrafish Maintenance

Zebrafish (*Danio rerio*) were kept under standard housing conditions as previously described (Brand et al., 2002). All fish used in this study were adults, both males and females, and 6 to 12 months of age. Transgenic fish used are: *Tg(−5.5opn1sw1:EGFP)^kj9^* (Takechi et al., 2003); *Tg(mpeg1:mCherry)* (Ellett et al., 2011); *Tg(mpo:GFP)* (Renshaw et al., 2016); *Tg(gfap:NLS-GFP)* (described here); *Tg(gfap:NLS-mCherry)* (Lange et al., 2020), *Tg(lck:NLS-mCherry)^sd31^* (Butko et al., 2015), *Tg(NF-kB:GFP)* (Kanther et al., 2011), Δirf8^st95^ (Shiau et al., 2015).

### Generation of Tg(gfap:nls-GFP)

The *Tg(gfap:NLS-GFP)* reporter line with its NLS-tag shows highly nuclear localization of GFP, but also some leaky cytoplasmic expression in Müller glial cells, and thus serves as an ideal marker for MGCs in our experiments. To generate this line, the GFP reporter was PCR amplified and flanked by restriction sites using GFP-for (atatGGCCGGCCgccaccatggctccaaagaagaagcgtaaggt) and GFP-rev (ggtgtgcatgttttgacgttgatggc) primers. By PCR, the nuclear localization sequence (NLS) was added as a 5’ overhang to the GFP. The PCR product was subcloned into the p2.1 Topo Vector. Next, Topo vector with the reporters and the *pTol(gfap:mcherry-T2A-CreERT2)* construct (SH and MB, unpublished) were digested using the enzymes Asc and FseI and ligated to replace the mCherry-CreERT2 cassette with the NLS-reporter. For germline transformation, either linearized plasmid DNA or plasmid DNA with transposase mRNA were injected into fertilized eggs (F0) in E3 medium (Brand et al., 2002), raised to adulthood and crossed to AB wild-type fish as previously described (Kawakami et al., 2004).

### Diffuse light lesion

To ablate photoreceptor cells, diffuse light lesion was performed as described in Weber et al., 2013. Briefly, dark-adapted fish were transferred to a beaker containing 250 ml system water and exposed to the light of an EXFO X-Cite 120 W metal halide lamp (∼200,000 lux) for 30’. For recovery, fish were connected to the system and kept under standard light conditions.

### Drug treatment

Dexamethasone (Dex, Sigma-Aldrich, Germany) was diluted from a 25 mg/ml stock solution in methanol to 15 mg/l in sterilized system water. For Dex treatment, adult fish were kept individually in sterilized tanks. Control groups with the corresponding amount of methanol (MeOH) without Dex, both groups did not show retinal cell death (supplement S3). Solutions were renewed daily, and tanks were exchanged every 5 days. Fish were fed with brine shrimp 1h prior to solution exchange. Animals were pretreated for 10 days and respective conditions were maintained until sacrificed. Light lesions were performed in corresponding solutions.

### Tissue preparation and sectioning

For retinal flat mounts, eyes were removed and a small incision in the cornea was made using a sapphire blade scalpel (ZT215 W0.5mm A60°). Eyes were prefixed in 4% PFA in calcium-free ringer solution (PFAR) for 30’ with slow agitation. Cornea, sclera and the lens were removed in calcium-free ringer solution and the retinae were cut at four sides. Flat mounts were fixed in 4% PFAR at 4 C overnight with gentle agitation. For storage, samples were transferred to 100% methanol and stored at - 20°C.

For sections, the lens was removed and fish heads were fixed at 4 C in 4% PFA in 0.1 M phosphate buffer (PB). Following that, samples were decalcified and cryo-protected with 20% sucrose/20% EDTA in 0.1 M PB. Tissues were embedded in 7.5% gelatine/20% sucrose in 0,1% PB, stored at -80 °C and sectioned into 14 µm cryo-sections using a Microm HM560. Sections were stored at -20°C.

### *In situ* Hybridization

*In situ* hybridization on sections was performed as described in Ganz et al., 2015. Hybridization was carried out overnight at 62 °C. Probes were generated using *mmp9* (primer forward: CTTGGAGTCCTGGCGTTTCT; primer reverse: GCCCGTCCTTGAAGAAGTGA) and *her4.1* (Takke et al., 1999) as targets. For detection, NBT/BCIP or SIGMA*FAST*^TM^ Fast Red TR/Naphtanol AS-MX was used. For combination with immunohistochemistry, the primary antibody was incubated with anti-digoxigenin-AP. Secondary antibody was applied after staining development for 2h and washed with PBS with 0.3% Triton-X100 (PBSTx). Sections were mounted in glycerol.

### Immunohistochemistry & TUNEL

Sections were dried for 2 h at 50 °C and rehydrated with PBSTx. Following this, sections were incubated with primary antibody: anti-Zrf-1 ZIRC mouse IgG1 1:200 (Fausett & Goldman, 2006), anti-Zpr-3 ZIRC IgG1 1:200 (Zou et al., 2008), anti-GFP Abcam chicken 1:3000 (Leinninger et al., 2009), anti-DsRed Clontech rabbit 1:500 (Glass et al., 2005), anti-PCNA mouse IgG2a DACO PC10 1:500 (Grandel et al., 2006), anti-L-Plastin rabbit 1:5000, (Redd et al., 2006) and incubated overnight at 4 °C. Excess antibody was washed off using PBSTx followed by incubation with secondary antibody against the respective primary antibody host (Molecular Probes; Alexa 488, Alexa 555, Alexa 635; 1:750) containing 1 µg/ml 4′,6-Diamidin-2-phenylindol (DAPI). Antibody-solution was washed off and slides were mounted with glycerol.

To retrieve the PCNA antigen, sections were incubated for 8’ in 50 mM Tris buffer pH 8.0 at 99 °C followed by 10 min PBS prior to primary antibody incubation.

Modifications for retinal flat mounts are prolonged incubation times for primary and secondary antibody solution for 48 h respectively at 4 °C with slow agitation and extensive washing 10x 30’ after each antibody incubation. Flat mounts were mounted on slides in glycerol.

TUNEL (TdT-mediated dUTP-biotin nick end labeling) assays were performed on sections using ApopTag® Red in Situ Apoptosis Detection Kit (Merk) according to manufacturer’s instructions.

### EdU labeling & detection

To trace proliferating cells, EdU (5-ethynyl-2′-deoxyuridine) pulses were given intraperitoneally. Fish were anesthetized in 0.024% Tricaine and EdU was injected intraperitoneally (20 µl of 2.5 mg/ml EdU in PBS per pulse). Detection was performed on sections using “Click-iT® Plus EdU Alexa Fluor® 555 Imaging Kit” (Thermo Fischer Scientific) according to manufacturer’s instructions.

### Intravitreal injections

Fish were anesthetized in 0.024 % tricaine and covered with tricaine-moistened paper towel, leaving the eye exposed. The outer cornea was removed. Subsequently, a small incision in the cornea was made using a sapphire blade scalpel (ZT215 W0.5mm A60°). A Hamilton syringe equipped with a 33 gauge, blunt-end needle was inserted and 0,5 µl of solution (flagellin from *S. typhimurium*, 0,1 µg/µl, Sigma; zymosan from *S. cerevisiae*, 0,2 µg/µl BioParticles; Human M-CSF, 50 ng/µl, PreproTech; Human Il-34, 50 ng/µl, PreproTech; PBS) was injected into the vitreous of the eye without touching the retina. The incision was covered with Histoacryl® (Braun) to avoid leakage.

### Image acquisition

Images were acquired using a Zeiss Axio Imager, equipped with ApoTome. EC Plan-Neofluar 5x/0.16, Plan-Apochromat 20x/0.8 and LD C-Apochromat 40x/1.1 W Korr UV VIS IR objectives were used for magnification. For detection either a monochromatic Axiocam HR (1388×1040 pixels, 6.45*6.45) for fluorescence or a polychromatic Axiocam MR Rev3 (1388×1040 pixels, 6.45*6.45) for bright-field imaging was used. Sequential image acquisition was used in co-stained samples with multiple fluorophores. Images were acquired in AxioVision Rel. 4.8 and processed using Fiji (Schindelin et al., 2012). Figure panels were assembled in Adobe Photoshop CS6.

### Cell counting and statistical analysis

For cell quantification, retinal sections with a thickness of 14 µm were used. For single fluorophore cell quantification, the first 3-5 consecutive retinal sections rostral to the optic nerve were selected and manually counted. For normalization, the length of the retina was determined for every section by measuring a snapshot of the retina at the level of the inner nuclear layer. For quantification of multiple fluorophores, 20x images were acquired. Cells were quantified using the cell counter tool in Fiji. Length measurements were applied on the same images for normalization. For statistical analyses, ≥3 fish were used for calculations. Counted cell numbers were normalized to the respective parameter (reference cell population or length) and the counts per fish were averaged. The average number for every fish was used as an analytical unit. For statistical analysis, Graph Pad Prism was used to determine p-values with a one-way ANOVA test with Tukey’s post hoc analysis. For unpaired two-tailed student’s T-Test, Excel was used. Significance levels are displayed as: not significant = p>0.05, * = p≤0.5, ** = p≤0.01, *** = p≤0.001. Graphs shown in bar charts were created using Excel, error bars represent standard error of the mean.

## Acknowledgments

We are very grateful to Marika Fischer, Jitka Michling, Daniela Mögel, and Claudia Meyer who provided outstanding fish care, and Dr. Judith Konantz for managing the fish facility. We also thank Michaela Geffarth and Nathalie Franke for excellent technical support and John F Rawls for sharing the *Tg(NF-kB:GFP)* reporter line. Further, we thank past and present members of the Brand lab for many interesting discussions, and Dr. Thomas Quail, Dr. Judith Konantz, Dr. Christian Lange, Prof. Dr. Thomas Becker, Prof. Dr. Catherina Becker, Prof. Dr. Marius Ader, Ecem Çayıroğlu and Prof. Dr. Susanne Koch for comments on earlier versions of this manuscript.

