## Supplementary Figures for "Successful regeneration of the adult zebrafish retina is dependent on inflammatory signaling"

**Supplement Figures**

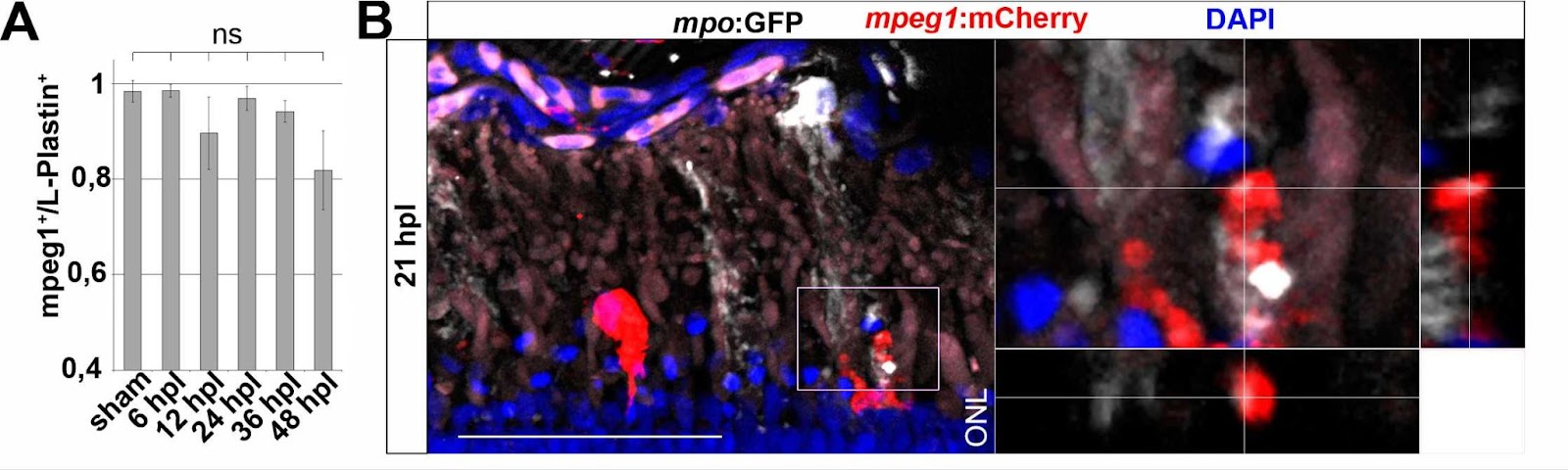

**Figure S1: Neutrophils as a part of *Tg(mpeg1:mCherry)-*negative leukocytes are attracted by lesion.** (A) Quantification of Tg(*mpeg1*:mCherry) and L-Plastin double positive cells in the context of lesion. The amount of double positive cells decreases, however not significantly. (B) Tg(*mpo*:GFP)^+^ neutrophils spread GFP positive matrixes at the side of injury. These structures are in close proximity to *Tg(mpeg1:mCherry)* positive monocytes. Scale bar=50µm; ONL=outer nuclear layer.

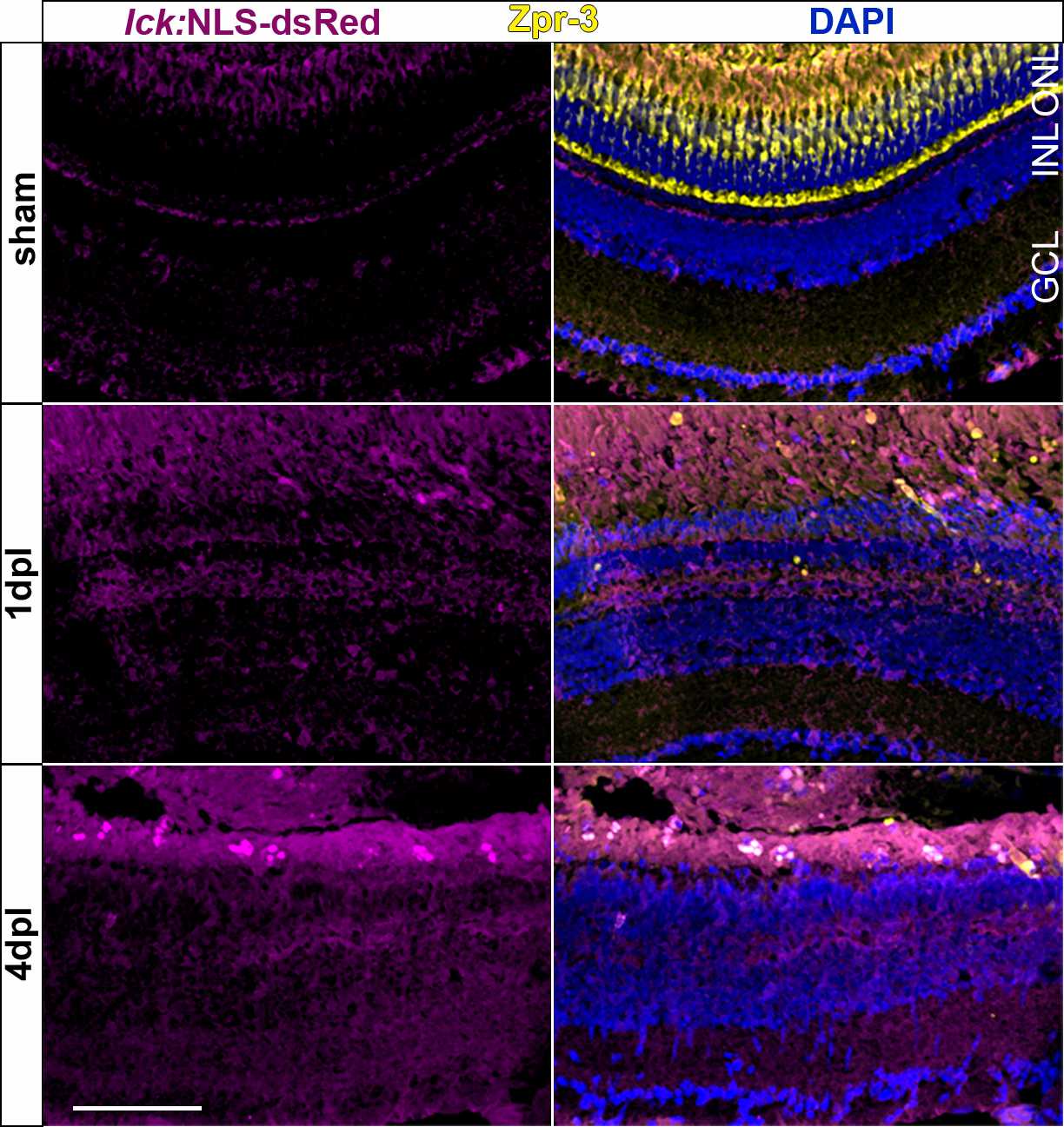

**Figure S2: *Tg(lck:NLS-dsRed)^+^* T cells do not respond to retinal injury.** No *Tg(lck:NLS-dsRed)* T cell is detected upon injury. Photoreceptor ablation is indicated by the lack of Zpr-3 positive red-/green-double cones at 1 and 4 days post lesion (dpl). Scale bar=50µm. ONL=outer nuclear layer; INL=inner nuclear layer; GCL=ganglion cell layer.

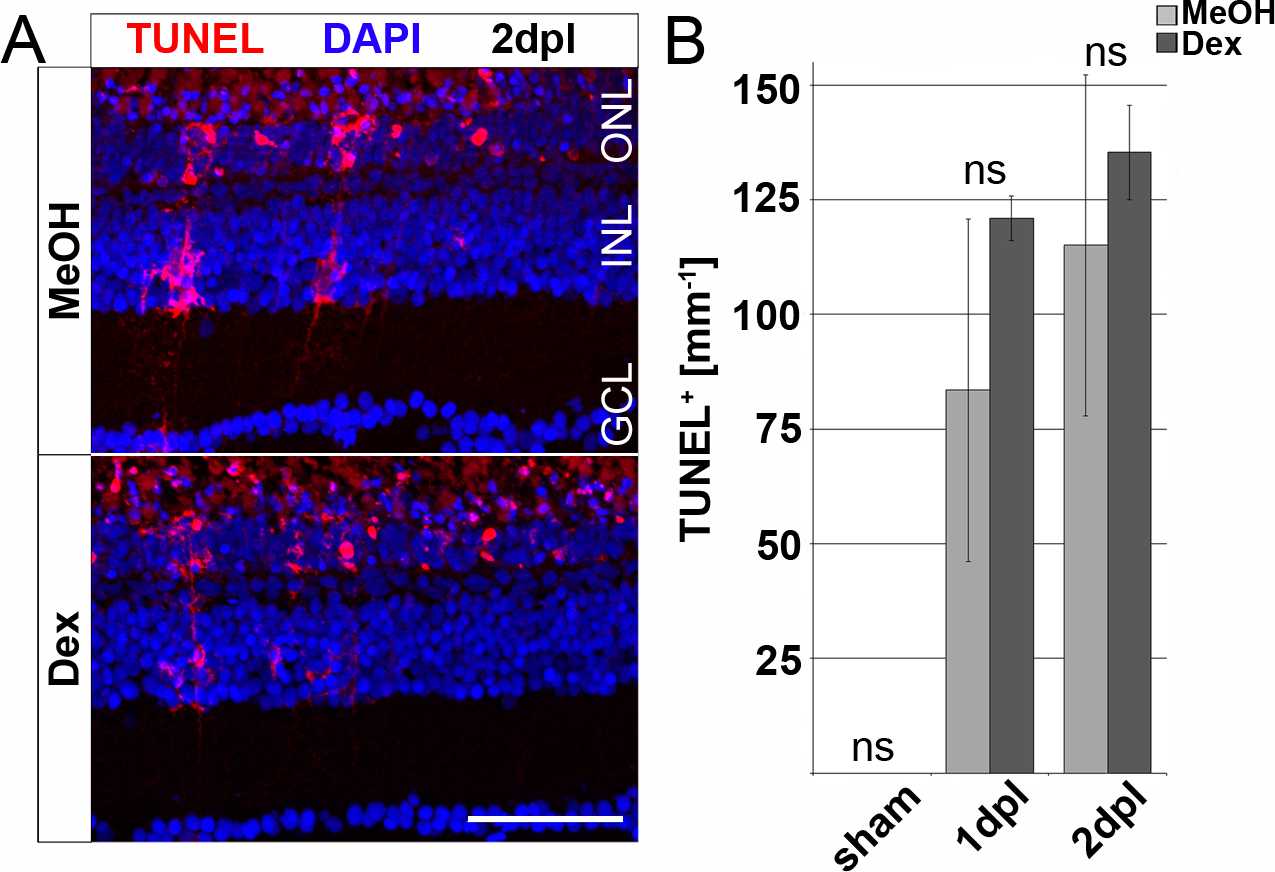

**Figure S3: Dexamethasone treatment has no influence on cell death induced by phototoxic ablation**. (A) TUNEL^+^ nuclei are detectable in the ONL of Dex and MeOH treated samples. (B) Quantification of TUNEL^+^ cells in sham, 1 dpl and 2dpl retina in Dex-treated eyes in comparison to vehicle control show no difference. Scale bar= 50 µm, Error bars indicate standard error; n.s. = p > 0,05; n=3; two-tailed student’s T-Test; ONL=outer nuclear layer; INL=inner nuclear layer; GCL=ganglion cell layer.

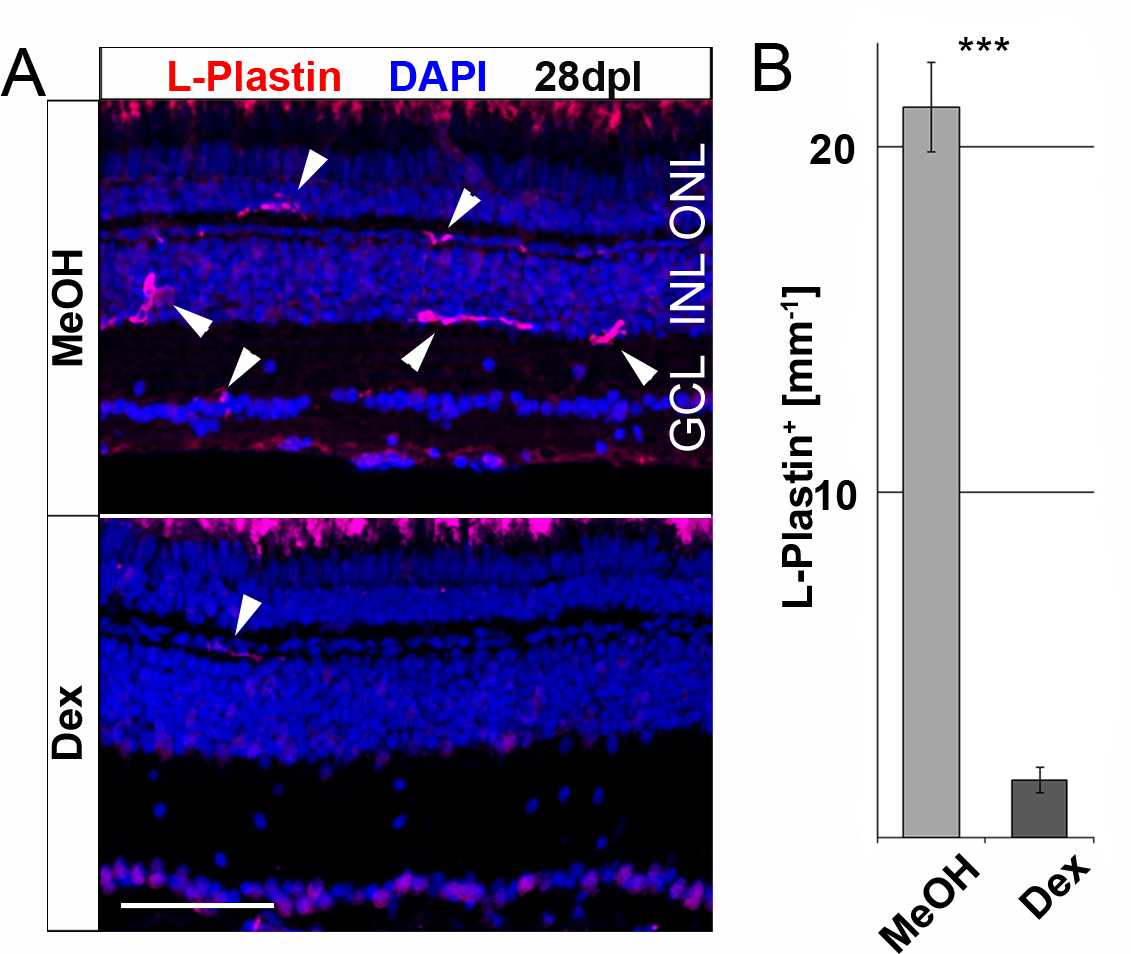

**Figure S4: Long-term dexamethasone treatment leads to a strong reduction of retinal leukocytes.** (A) After 48 days of continuous Dex-treatment L-Plastin^+^ cells can rarely be detected in Dex samples in comparison to MeOH control. (B) Quantification of L-Plastin^+^ cells in the retina of MeOH and Dex-treated fish reveals a strong decrease of L-Plastin^+^ cells. Scale bar= 50 µm, Error bars indicate standard error; *** = p < 0,001; n=6; two-tailed student's T-Test; ONL=outer nuclear layer; INL=inner nuclear layer; GCL=ganglion cell layer.
